# Mechanistic insights into ligand dissociation from the SARS-CoV-2 spike glycoprotein

**DOI:** 10.1101/2023.11.29.569184

**Authors:** Timothy Hasse, Esra Mantei, Rezvan Shahoei, Shristi Pawnikar, Jinan Wang, Yinglong Miao, Yu-ming M. Huang

## Abstract

The COVID-19 pandemic, driven by the severe acute respiratory syndrome coronavirus 2 (SARS-CoV-2), has spurred an urgent need for effective therapeutic interventions. The spike glycoprotein of the SARS-CoV-2 is crucial for infiltrating host cells, rendering it a key candidate for drug development. By interacting with the human angiotensin-converting enzyme 2 (ACE2) receptor, the spike initiates the infection of SARS-CoV-2. Linoleate is known to bind the spike glycoprotein, subsequently reducing its interaction with ACE2. However, the detailed kinetics underlying the protein-ligand interaction remains unclear. In this study, we characterized the pathways of ligand dissociation and the conformational changes associated with the spike glycoprotein by using ligand Gaussian accelerated molecular dynamics (LiGaMD). Our simulations resulted in eight complete ligand dissociation trajectories, unveiling two distinct ligand unbinding pathways. The preference between these two pathways depends on the gate distance between two α-helices in the receptor binding domain (RBD) and the position of the N-linked glycan at N343. Our study also highlights the essential contributions of K417, N121 glycan, and N165 glycan in ligand unbinding, which are equally crucial in enhancing spike-ACE2 binding. We suggest that the presence of the ligand influences the motions of these residues and glycans, consequently reducing accessibility for spike-ACE2 binding. These findings enhance our understanding of ligand dissociation from the spike glycoprotein and offer significant implications for drug design strategies in the battle against COVID-19.

## Introduction

The ongoing COVID-19 pandemic, caused by the severe acute respiratory syndrome coronavirus 2 (SARS-CoV-2), has had a significant global impact on health and economies, highlighting the urgent need for effective therapeutic interventions^1,2^. The spike glycoprotein of SARS-CoV-2 plays a critical role in viral entry into host cells, facilitated by its interaction with the human angiotensin-converting enzyme 2 (ACE2) receptor^3,4^. Consequently, the development of small molecules capable of disrupting the spike-ACE2 interaction emerges as a promising avenue for antiviral treatments^5,6^. In the past, researchers primarily focused on assessing protein-ligand binding affinity by measuring the equilibrium dissociation constant (*K_D_*) or half-maximal inhibitory concentration (*IC*_50_)^7,8^. However, recent findings have emphasized the importance of considering the kinetics of ligand binding and unbinding, characterized by the ligand association rate constant (*K_on_*) and dissociation rate constant (*K_off_*), respectively^9,10^. By taking into account these kinetic properties, valuable insights can be gained regarding the dynamic nature of protein-ligand interactions and their impact on drug efficacy. Given that similar intermediates may occur in both ligand binding and unbinding processes^11^, our study is dedicated to unraveling the intricate process of ligand dissociation. Through this investigation, we aim to provide valuable insights into the kinetics of ligand diffusion and the intrinsic dynamical behavior observed during ligand unbinding processes.

The spike glycoprotein is composed of three identical chains (Figure 1A and 1B), each consisting of two functional subunits: the S1 subunit, which contains the receptor-binding domain (RBD), and the S2 subunit (Figure S1A), which facilitates membrane fusion^12–14^. The RBD binds to the ACE2 receptor on host cells, particularly through the receptor-binding motif (RBM), initiating the viral entry process^15^. The spike glycoprotein exists in two primary conformations: a “closed” state, characterized by all RBDs being in a “down” conformation, rendering them inaccessible for ACE2 binding; and an “open” state, where at least one RBD is in an “up” conformation, facilitating exposure and enabling binding to ACE2^16,17^.

**Figure 1.**
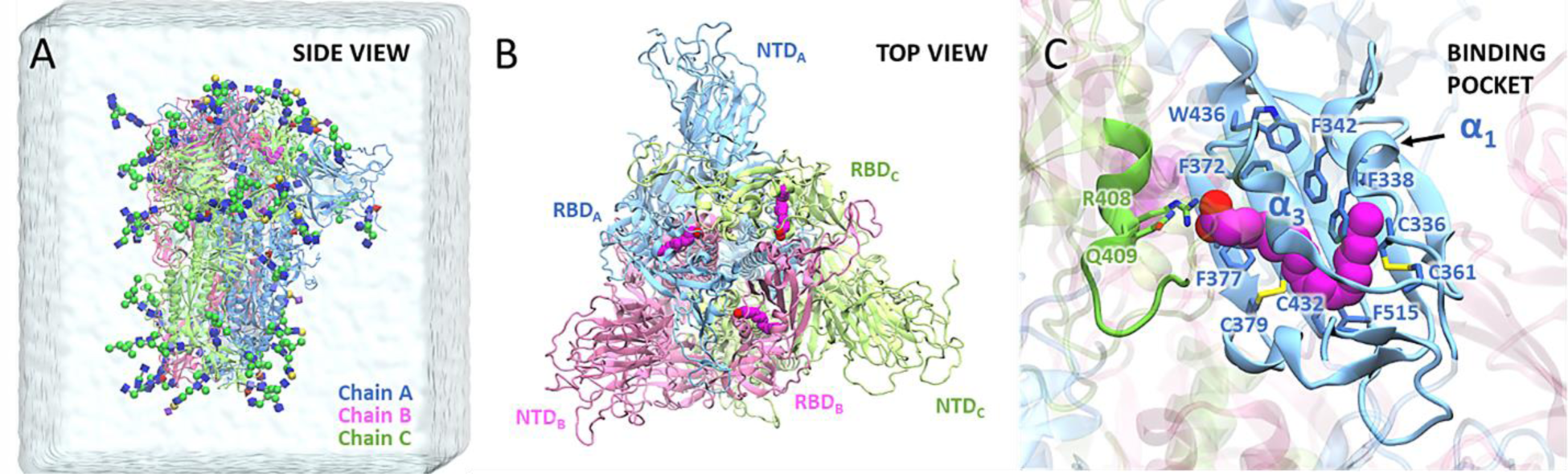
Structure of spike glycoprotein with bound linoleate (LA) molecules. (A) Side view of the spike trimer, comprising Chain A (blue), Chain B (pink), and Chain C (green). The structure is shown in the simulation water box with glycans in the 3D-SNFG cartoon representation. (B) Top view of the spike trimer, highlighting the receptor-binding domain (RBD) and N-terminal domain (NTD). Each RBD contains a bound LA molecule (magenta), with the corresponding subunit indicated as A, B, or C, noted as subscripts. (C) Close-up view of the free fatty acid (FFA) binding pocket within the RBD, illustrating the specific interaction of LA within the pocket.

To design inhibitors that can disrupt the spike-ACE2 interactions, extensive investigations have explored the binding affinities of the SARS-CoV-2 spike glycoprotein with various compounds, such as antibodies^18^, peptides^18,19^, carbohydrates^20^, and polyunsaturated fatty acids^21^. Among these spike binders, linoleate (LA), a free fatty acid (FFA) known for its crucial role in inflammation and immune modulation^22,23^, has demonstrated remarkable potency in blocking the entry of SARS-CoV-2^24^. The structure of the spike-LA complex has been revealed through cryo-electron microscopy, uncovering the binding of LA to a hydrophobic pocket within the RBD^24^ (Figure 1C). The FFA binding pocket, enclosed by 6 α-helices and 13 β-sheets^25^ (Figure S1B), positions at the interface of two RBDs from neighboring chains. The LA hydrophobic tail interacts with nonpolar residues of the α1 and α3 helices and the β-sheet on the primary RBD, while the LA hydrophilic head interacts with polar residues on the neighboring RBD (Figure 1C). Additionally, experimental data for *K_D_*, as well as the *K_on_* and *K_off_* of LA, have been reported^24^. The availability of structural information along with thermodynamic and kinetic properties makes LA an excellent candidate for diffusion mechanism studies that can serve as a model for other potential spike binders.

Computational methods are crucial in spike glycoprotein studies, allowing for in-depth exploration of spike structure, dynamics, and interactions. These methods provide valuable insights that complement experimental data and could greatly contribute to the discovery of potential therapeutics and interventions. For example, *Casalino et al.*^26^ investigated the multifaceted roles of glycans in spike viral entry and immune recognition, *Kapoor et al.*^27^ highlighted the impact of posttranslational modifications on spike glycoprotein interactions with host cell receptors, and *Wang et al.*^5^ utilized *in silico* techniques to explore the potential of small molecules targeting the conserved FFA binding pocket of the spike glycoprotein as a starting point for broad-spectrum COVID-19 intervention treatments. Despite advancements in spike glycoprotein studies, a detailed understanding of ligand binding and unbinding mechanisms remains limited. To address the knowledge gap, we employ ligand Gaussian accelerated molecular dynamics (LiGaMD)^28^ to investigate the kinetics of ligand dissociation from the spike glycoprotein. LiGaMD achieves enhanced sampling of ligand unbinding by selectively adding a harmonic boost potential to the ligand nonbonded interaction energy, enabling accelerated observations of the spike-ligand dissociation process without predefined reaction coordinates^28^.

In this study, we conducted all-atom LiGaMD simulations on spike glycoprotein systems with bound LA molecules to characterize the pathways of ligand dissociation and the conformational changes associated with the spike glycoprotein. By analyzing the critical interactions and key residues present in each intermediate state, as well as studying the transitions occurring along these pathways, we identified distinct pathways for LA unbinding. We also investigated the corresponding protein conformational changes and examined the influence of glycans on ligand dissociation. Our study sheds light on the kinetic process of ligand unbinding, further enhancing the understanding of the underlying mechanisms that govern ligand interactions, facilitating optimized strategies for drug design.

## Methods

### Model systems

The SARS-CoV-2 spike protein complex, consisting of three bound LA molecules, was obtained from the Protein Data Bank (PDB) under the code 6ZB5^24^. This protein complex exhibited a closed conformation with C3 symmetry. The missing loops were built based on the structure reported by *Casalino et al*.^26^ We constructed spike protein systems with varying ligand configurations, including those with one, two, or no ligands present, achieved by manually removing LA molecules from the FFA binding pocket. The protonation states of protein sidechains were determined using the PROPKA program through the PDB2PQR server^29^. Additionally, glycan attachments onto the protein were built using the CHARMM-GUI online server^30,31^, following the glycan positioning established in prior research by *Casalino et al.*^26^ (Table S1).

### Simulation protocol

The AMBER20 simulation package^32^ was employed for system preparation and enhanced sampling, The AMBER ff19SB force field^33^ was utilized for the protein, while GLYCAM_06j^34^ and GAFF2^35^ were used for carbohydrates and ligands, respectively. The systems underwent a comprehensive three-stage minimization process. This process involved 5,000 steps of hydrogen minimization, followed by 25,000 steps of hydrogen, sidechain, ligand, and glycan minimization. The final step included 25,000 steps of whole system minimization. Subsequently, the systems were solvated using the TIP3P water model^36^, extending to 20 Å from the protein surface. The systems were neutralized and Na^+^ and Cl^-^ ions were introduced at a concentration of 0.15 M to match extracellular NaCl concentration. A secondary round of minimization was conducted, with 5,000 steps dedicated to minimization of water molecules, followed by 25,000 steps of whole system minimization. Following this, the system underwent equilibration through a series of steps. Initial solvent equilibration was performed in the NPT ensemble at 310 K for 20 ps while restraining the protein. Subsequent equilibrations were performed in the NVT ensemble at 50, 100, 150, 200, 250, and 310 K without restraints. A final conventional MD simulation of 20 ns at 310 K ensured the system convergence to the appropriate thermodynamic internal energy before commencing LiGaMD simulations.

LiGaMD is an enhanced sampling technique designed to efficiently explore ligand unbinding pathways by adding a selective boost potential energy to the nonbonded interactions between the ligand and receptor^28^. The specifics of LiGaMD are elaborated in Text S1. Our LiGaMD simulations comprised an 84 ns preparation phase, followed by a production run of 500 ns or until successful ligand dissociation, whichever happened first. The preparation run commenced with a 4 ns conventional MD stage at 310 K, used to gather potential energy statistics necessary for calculating the harmonic force constant and threshold energy parameters essential for the boost potential evaluation. Next, a 1 ns LiGaMD simulation applied the boost potential without parameter updates. Subsequently, during an 80 ns LiGaMD simulation, the boost parameters were iteratively updated every 1 ns to determine new boost potentials. During the production run, the boost potential remained constant across the entire simulation, applied to both the ligand nonbonded potential energy and the potential energy of the remaining system components for optimal acceleration. The simulations adhered to an upper bound energy limit, maintaining the upper limit of the standard deviation of the ligand nonbonded boost potential at 6.0 kcal/mol. The upper limit for the boost potential of other energy terms was set in the range of 60-150 kcal/mol to enhance the success of ligand diffusion. The details of LiGaMD parameters can be found in Text S2. All LiGaMD simulations employed Langevin dynamics^37^ with a collision frequency of 5 ps^-1^. A 9 Å cutoff was applied, and the particle mesh Ewald summation^38^ was enabled. Hydrogen-containing bonds were restrained using the SHAKE algorithm^39^, and the simulation time step was set to 2 fs. Trajectories were recorded every 10 ps for subsequent analysis.

### Simulation analysis

The analysis of simulation trajectories, encompassing heavy-atom root-mean-square deviation (RMSD), root-mean-square fluctuation (RMSF), distance measurement, and dihedral calculations, was executed using VMD^40^ and CPPTRAJ^41^ tools. The RMSD of LA relative to the initial equilibrated structure was computed to elucidate the ligand movement throughout the dissociation trajectory. Additional RMSD calculations were carried out for the complete spike protein as well as for the residues within the RBDs. The analysis also involved the calculation of the RMSF to examine the flexibility and rigidity of individual spike protein residues. Two-dimensional potential of mean force (PMF) profiles were derived from the LiGaMD trajectories without energetic reweighting, utilizing the PyReweighting toolkit^42^. The PMF, which serves as an expression of free energy along reaction coordinates, was computed using the equation: *F*(*A*) = −*k*_*B*_*T* ln *P*(*A*), where *F*(*A*) represents the free energy along the reaction coordinate *A*, *k*_*B*_ is the Boltzmann constant, *T* is temperature, and *P*(*A*) denotes the probability distribution of *A*. The bin size for both ligand RMSD and atomic distance was set at 0.5 Å.

## Results

In this study, we aim to unravel the mechanism of LA unbinding from the spike-LA complex structure by conducting LiGaMD simulations. The complex structure initially contains three LA molecules bound to the spike glycoprotein. To explore the intermediate interactions and different binding pockets during metastates, we performed LiGaMD simulations on seven holo spike systems, each containing varying numbers of LA molecules (one, two, or three). Since the current version of LiGaMD allows selective boosting of only one bound ligand in the simulation, we designed twelve simulation systems to encompass all possible scenarios. These systems included LA bound to different RBD subunits and were denoted as 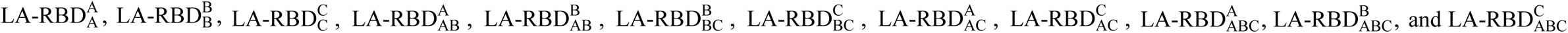. The subscripts indicate the RBD subunit(s) that have LA bound molecules at the start of the simulations, while the superscripts represent the RBD subunit that the boosted ligand was bound to. To achieve successful ligand dissociations, we gradually increased the boost potential, starting from a low value and incrementally raising it until the ligand dissociation event was observed. Notably, although the spike protein exhibits trimeric symmetry with identical protomers, our models incorporated specific glycan types and positions, which were found to differ between each chain based on previous work^26,43^. This distinction resulted in unique environments for the ligands bound to each chain.

### Two distinct LA dissociation pathways captured by LiGaMD simulations

The ligand movement was evaluated by calculating the heavy-atom RMSD of the LA molecule relative to its initial equilibrated structure. Complete dissociation of the ligand from the binding pocket was defined as when the ligand RMSD values exceeded 30 Å. Our LiGaMD simulations yielded a total of eight successful dissociation events, with dissociation times ranging from approximately 100 to 400 ns (Table 1 and Figure 2A). Among these eight dissociation events, four originated from the system with a single LA bound initially to RBD_A_. Two dissociation events occurred from LA bound to RBD_B_, involving one in a system with a single bound ligand and the other with three bound ligands. The remaining two dissociation events involved the unbinding of LA from RBD_C_ in systems with two and three bound ligands (Table 1).

**Figure 2.**
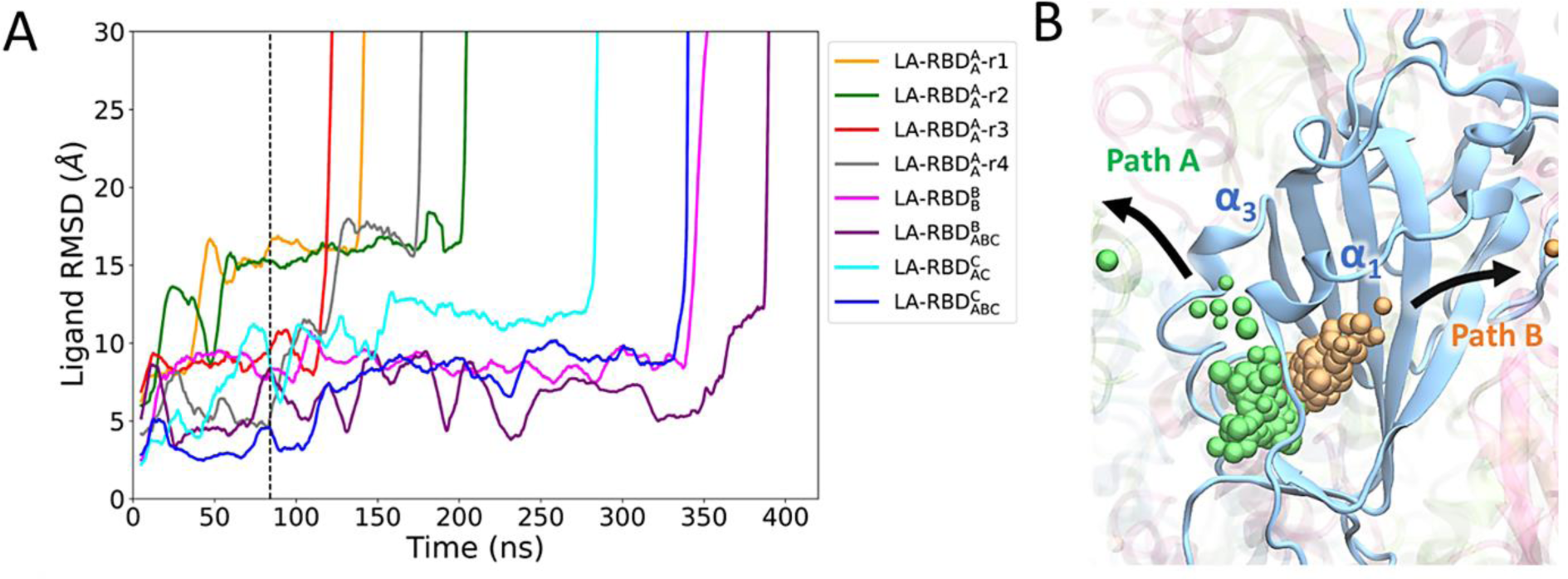
Linoleate (LA) dissociation pathways. (A) The RMSD of LA relative to the initial equilibrated structure for each successful dissociation simulation. A successful dissociation is defined by a ligand RMSD greater than 30 Å. The simulation time before and after the dashed line indicates the GaMD preparation and production run, respectively. (B) Representative trajectories depicting two distinct dissociation pathways: Path A (green) and Path B (orange). The ligand trajectory is shown by individual beads, representing the center of mass coordinates of LA over the simulation time.

**Table 1.** List of successful spike-LA dissociation simulations. The subscript and superscript of RBD indicate the domains that contain bound and boosted LAs, respectively. σ_L,nb_ and σ_D_ represent the user-defined upper limit of the standard deviation of the boost potential added to the nonbonded ligand potential terms and to the rest of the potential terms, respectively (details are in Text S1).

| System | Bound Ligands | Boosted Ligand | $\sigma_{L,nb}$<br>(kcal/mol) | $\sigma_D$<br>(kcal/mol) | Simulation Time | Dissociation Time | Dissociation Pathway |
| --- | --- | --- | --- | --- | --- | --- | --- |
| LA-RBD<sub>A</sub><sup>A</sup>-r1 | RBD<sub>A</sub> | RBD<sup>A</sup> | 6.0 | 60.0 | 160.0 ns | 143.4 ns | B |
| LA-RBD<sub>A</sub><sup>A</sup>-r2 | RBD<sub>A</sub> | RBD<sup>A</sup> | 6.0 | 150.0 | 200.0 ns | 184.4 ns | B |
| LA-RBD<sub>A</sub><sup>A</sup>-r3 | RBD<sub>A</sub> | RBD<sup>A</sup> | 6.0 | 150.0 | 140.0 ns | 119.7 ns | A |
| LA-RBD<sub>A</sub><sup>A</sup>-r4 | RBD<sub>A</sub> | RBD<sup>A</sup> | 6.0 | 150.0 | 200.0 ns | 178.2 ns | B |
| LA-RBD<sub>B</sub><sup>B</sup> | RBD<sub>B</sub> | RBD<sup>B</sup> | 6.0 | 150.0 | 360.0 ns | 348.7 ns | A |
| LA-RBD<sub>ABC</sub><sup>B</sup> | RBD<sub>ABC</sub> | RBD<sup>B</sup> | 6.0 | 120.0 | 420.0 ns | 392.0 ns | A |
| LA-RBD<sub>AC</sub><sup>C</sup> | RBD<sub>AC</sub> | RBD<sup>C</sup> | 6.0 | 150.0 | 300.0 ns | 287.0 ns | B |
| LA-RBD<sub>ABC</sub><sup>C</sup> | RBD<sub>ABC</sub> | RBD<sup>C</sup> | 6.0 | 60.0 | 360.0 ns | 343.1 ns | A |

During these dissociations, two distinct unbinding pathways, referred to Path A and Path B (Figure 2B), were identified. Path A entails the movement of LA between the α1 and α3 helices, whereas Path B involves LA exiting the pocket through the region between the α1 helix and the β-sheet. Notably, our findings did not indicate any evident correlation between the unbinding pathway and the initial binding pocket, the initial number of bound ligands in the system, or the applied boost potential.

Our simulations started with the LA molecule fully bound in the FFA binding pocket. The polar head of LA predominantly interacts with polar residues R408, Q409, and K417 on the neighboring RBD loop, forming a polar anchor, while the hydrophobic tail is positioned between the α1 and α3 helices as well as the β-sheet of the primary RBD, effectively locking LA in place (Figure S2A). Following the initial bound state, LA undergoes a flipping motion, reorienting itself within the binding pocket. This motion causes the polar head group of LA to face away from the neighboring RBD and eventually exit the binding pocket, following either Path A or Path B. (Figure S2B). The ligand flipping motion typically occurs within the initial 20 ns, and the subsequent dissociation of the polar interaction transpires within the first 60 ns. During LA dissociation, its polar head extends out of the binding pocket and positions the ligand between α1 and α3 for Path A, or between α1 and the β-sheet for Path B. Concurrently, the hydrocarbon tail gradually moves away from the surrounding hydrophobic residues (Figure S2C and S2D). LA may rebind to the pocket due to the interplay between solvent attraction to the LA polar head and the interaction between the fatty acid tail and hydrophobic residues of the binding pocket. This delicate balance between solvent and hydrophobic interactions plays a crucial role in determining whether LA remains in the bound state or dissociates from the pocket. Ultimately, regardless of whether LA follows Path A or Path B, both the polar head and the nonpolar tail of LA become fully exposed to the solvent (Figure S2E and S2F). It is worth noting that all unboosted ligands show minimal fluctuations and consistently remain fully bound throughout the simulations, resembling their initial conformation. This highlights their stability within the binding pocket during the simulations.

### Conformational flexibility of the spike glycoprotein during LA dissociation

During the dissociation of LA from the complex, both the ligand and the spike glycoprotein experience significant conformational changes and dynamical fluctuations. We conducted an analysis to assess the conformational flexibility of the spike glycoprotein and the regions involving the RBDs during the ligand dissociation process. To examine the overall motion of the spike glycoprotein, we calculated the RMSD of the entire spike throughout the simulations (Figure S3). The RMSD values generally increased as the ligand dissociation progressed. Initially, the change of RMSD was around 4 Å, but it typically rose above 6 Å and occasionally approached 8 Å, indicating deviations from the initial state and highlighting the impact of ligand dissociation on the overall protein structure. Next, we focused on the RMSD of the RBDs (Figure S4), as these regions directly participate in ligand unbinding. Throughout the ligand dissociation trajectories, the RMSD of each RBD exhibited fluctuations. Specifically, when LA transitioned from the initial bound state to either Path A or Path B, the RMSD of the corresponding RBD increased, indicating conformational changes associated with the ligand dissociation. Interestingly, in cases where the LA polar head rebound to the polar anchor residues R408, Q409, and K417, the RMSD of the RBD decreased rapidly within a short period of time. This dynamic behavior of the RBDs suggests structural changes accompanying the ligand dissociation process, significantly influenced by the ligand interaction with nearby residues.

We conducted RMSF analysis to explore the dynamic fluctuations of individual residues within the RBDs during LA dissociation (Figure S5). Our investigation focused on the RMSF profiles of the RBDs containing a boost ligand in three distinct states: 1) the fully bound state, 2) during traversal along either Path A or Path B, and 3) following complete dissociation of LA from the RBD. Figure S5 illustrates that throughout the dissociation trajectory, both the α1 and α3 helices exhibit significant fluctuations, while the β-strands shown in Figure S2 generally display more stability. As anticipated, the dissociation of the ligand along either Path A or Path B results in a significant enhancement in the fluctuation of all residues. After LA fully dissociated, the RMSF values reduce to the level that is similar to those observed in the fully bound state. Additionally, the RBM region, which houses crucial residues interacting with the ACE2 receptor during viral entry^15^, was the most flexible region during LA dissociation.

Given the importance of α1 and α3 during ligand dissociation, we conducted further investigations to explore their significance in the process of LA unbinding. The α1 and α3 helices of the RBD function as a gate that undergoes conformational changes to accommodate the movement of the ligand unbinding from the pocket. In Path A, the α1 and α3 helices move apart from each other, creating a space that enables the ligand to exit the pocket between them. (Figure 3A). On the other hand, Path B requires α1 to move closer to α3, permitting the ligand to leave through the pocket between α1 and the β-sheet (Figure 3B). To quantitatively analyze these structural changes and the gate controlled by α1 and α3, we measured the distance between residues E340 of α1 and A372 of α3. We defined the gate open and closed states by distances of 18.21 Å (PDB ID 6VXX^44^) and 13.31 Å (PDB ID 6ZB5^24^), respectively, according to the investigation of multiple spike structures from the PDB (Table S2). Throughout the ligand unbinding trajectory, the gate distance fluctuates between the open and closed states. Specifically, for ligands following Path A, the gate must open to a distance greater than 18.21 Å to enable the ligand to pass through (Figure 3C). Conversely, Path B does not necessitate the gate to fully close to 13.31 Å, although the gate distance below 15.0 Å was required for the ligand to exit the binding pocket (Figure 3D). These findings emphasize the dynamic nature of the gate and its role in facilitating the ligand dissociation process along both pathways.

**Figure 3.**
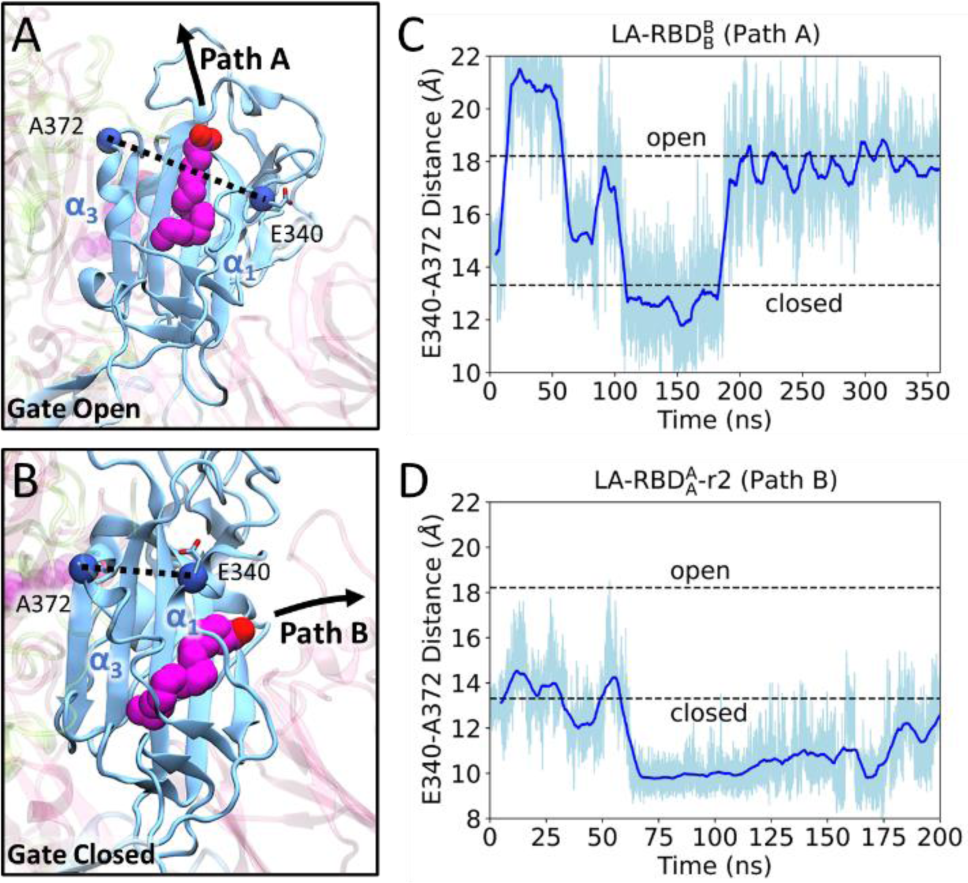
Dynamic opening and closing of the FFA binding pocket. (A) Ligand dissociation along Path A involves LA passing through the open gate formed by the α1 and α3 helices. The opening of the gate is measured by the distance between E340 on α1 and A372 on α3. (B) Ligand dissociation along Path B occurs when α1 and α3 move closer, resulting a closed gate and allowing the ligand to move between α1 and β-sheets. The distance between E340 and A372 for a Path A (C) and Path B (D) changes over the simulation time.

### Free energy profiles of LA unbinding from the spike glycoprotein

Utilizing the ligand RMSD and gate distance as reaction coordinates, we constructed a potential of mean force (PMF) diagram (Figure 4) to reveal the energetic change of LA dissociation. The PMF analysis revealed five local energy minimum states: the initial fully bound state and four intermediate states. During the ligand unbinding along Path A, the ligand transitions from the bound state to intermediate I1A, then to intermediate I2A, and ultimately becomes unbound. Conversely, for Path B, the ligand moves from bound to I1B, then to I2B, and finally becoming unbound.

**Figure 4.**
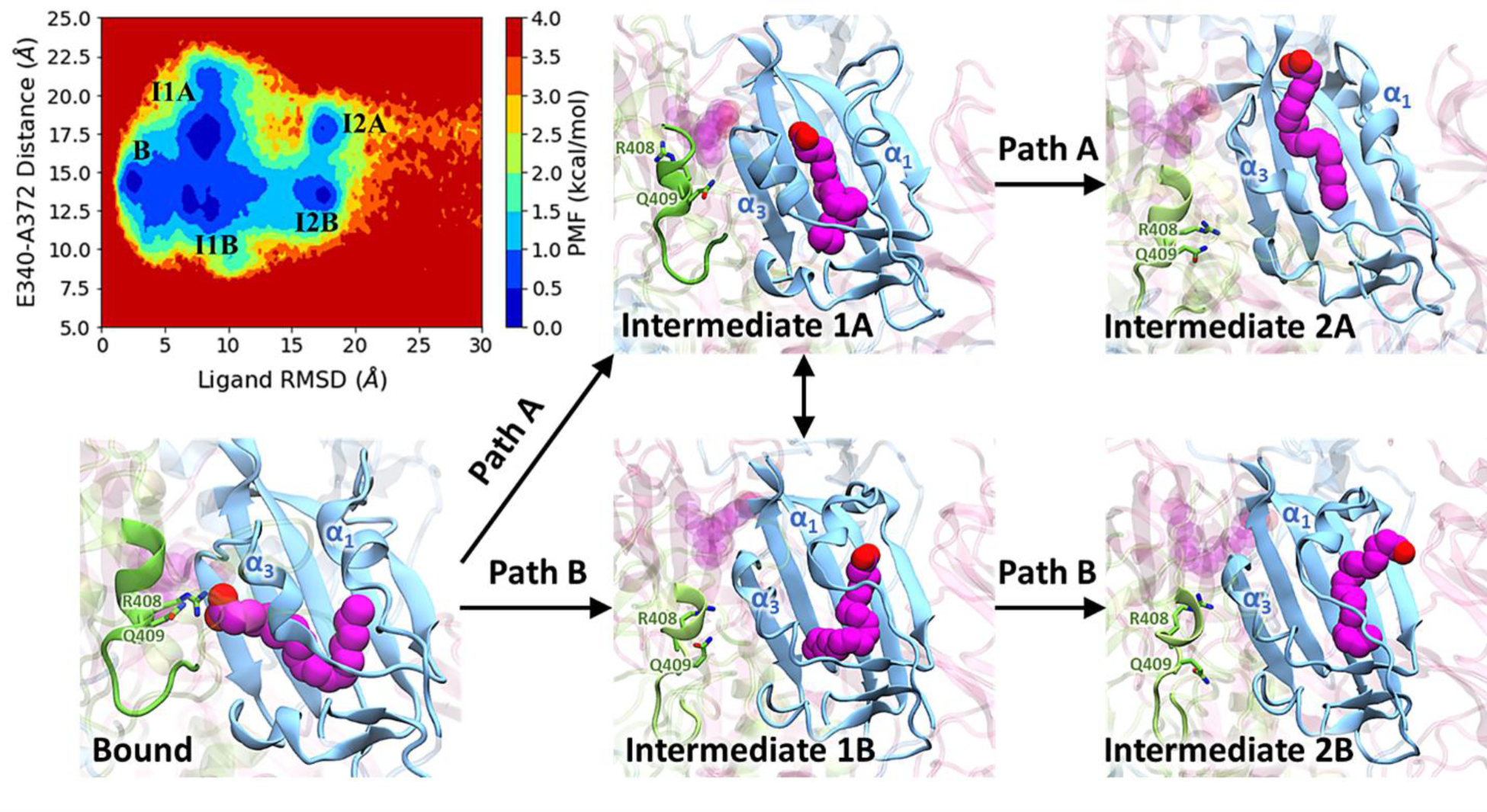
PMF profile of LA dissociation. The 2D PMF plot illustrates ligand RMSD and gate distance changes, indicating the intermediate conformations during LA unbinding processes. Arrows show the LA snapshots, starting in the fully bound state and moving along either Path A or Path B. A dual arrow indicates the transition of LA between dissociation paths at intermediate stages I1A and I1B.

Initially, in the ligand-bound state, LA is fully buried in the binding pocket formed by α1, α3, and β-sheet, while the gate distance is approximately 14.5 Å. As the ligand begins to dissociate, the polar interactions between LA and the residues of the RBD break, and the carboxyl headgroup of LA starts to interact with the surrounding water molecules outside the binding pocket, resulting in intermediate states I1A and I1B. In these states, the ligand can transition between intermediate states I1A and I1B, as the gate swings open and closed with the gate distance ranging from 12.5 Å to 17.5 Å. Moreover, the ligand can rebind to the binding pocket by forming hydrophobic interactions with the residues on the β-sheet close to α1 and α3. Subsequently, the ligand will continue to move farther away from the center of the pocket. Along Path A, the hydrophobic tail of LA is located between α1 and α3, forming nonpolar interactions with the α-helices, and the gate distance is around 17.5 Å, corresponding to intermediate state I2A. On the other hand, along Path B, LA is positioned between α1 and the β-sheet, with the gate distance closing to approximately 13.0 Å, denoted as intermediate state I2B. Once LA reaches either state I2A or I2B, it eventually dissociates from the FFA binding pocket. After dissociation, the gate distance exhibits fluctuations within the range of 10-20 Å, while the ligand RMSD can reach up to 200 Å, consistently remaining greater than 30 Å (Figure S6).

### N-Glycan at N343 modulates the LA unbinding pathways

The glycan at N343 located on the α1 helix plays a crucial role in determining the pathways of ligand unbinding. Our findings reveal a synchronous motion between the N343 glycan and the α1 helix. The dihedral angle shown in Figure 5C experiences significant changes as the ligand dissociates from the spike along different pathways. In Path A, the glycan rotates away from the pocket between α1 and α3 (Figure 5A), resulting in the fluctuation of dihedral angle between 200 and 280 degrees (Figure 5C). Conversely, during the dissociation of LA along Path B, the α1 helix swings towards α3, effectively closing the gate, while the glycan also moves towards α3, thereby hindering the pocket between α1 and α3 and impeding path A (Figure 5B). Throughout this trajectory, the dihedral angle fluctuates between 100 and 180 degrees (Figure 5D).

**Figure 5.**
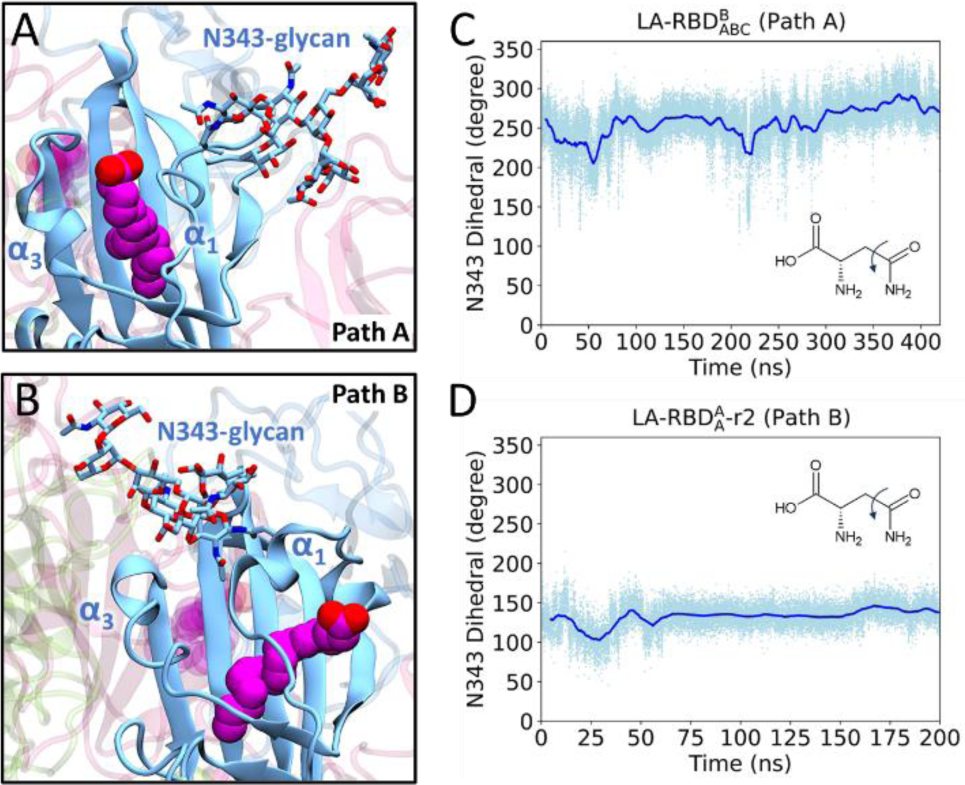
Dynamics of the glycan at N343. (A) In the case of dissociation along Path A, the N343 glycan extends into the solvent, moving away from the α1 and α3 helices. This allows LA to traverse through the open gate created by the α1 and α3 helices. (B) Conversely, during dissociation along Path B, the N343 glycan moves towards the α1 and α3 helices, restricting LA from progressing through Path A. Measurement of the dihedral angle of residue N343 is shown for Path A (C) and Path B (D), spanning the simulation duration.

Additionally, the orientation of the glycan at N343 also plays a role in influencing the dissociation of LA by engaging in interactions with other residues or glycans within the spike glycoprotein. Our analysis of LA unbinding trajectories revealed two predominant states of the N343 glycan. In one state, the glycan shifts toward the N-terminal domain (NTD), while in the other state, it moves closer to the RBM. When the glycan adopts the former orientation, its outward positioning exposes the region between α1 and α3, which potentially facilitates the movement of LA along Path A. In this configuration, the glycan extends further into the solvent and establishes interactions with two glycans bound to residues N122 and N165 (Figure S7A). Conversely, in the latter state, the glycan interacts with residues within the RBM, including K417, Y453, L455, F486, N487, Y489, Q493, Q498, N501, and Y505 (Figure S7B-D). This interaction pattern constrains the adjacent RBD conformation and enhances the accessibility of LA dissociation along Path B.

## Discussion

Understanding the intricacies of ligand dissociation processes remains paramount in the pursuit of designing novel inhibitors. Conventional MD simulations are often insufficient in exploring complete ligand dissociation events given that their typical sampling timescales fall within the range of nanoseconds to microseconds^45,46^. This timeframe is not able to fully capture unbinding processes, which usually extend from microseconds to seconds^47^. While several enhanced sampling methods, such as umbrella sampling^48,49^, metadynamics^50,51^, adaptive biasing force^52^, and steered MD^53^, are employed to investigate atomistic details of ligand unbinding, they commonly rely on predefined reaction coordinates. In this study, we applied LiGaMD^28^ to investigate ligand dissociation processes without introducing biased results from preselected reaction coordinates. Specifically, the method achieves acceleration by enhancing the nonbonded interactions within the ligand-binding site, enabling the sampling of elusive LA unbinding events^28^. Our findings uncover two distinct LA dissociation pathways and their corresponding intermediate states. Moreover, our analysis highlights the significance of the α1 and α3 gate and the N343 glycan in regulating the preferred LA unbinding routes.

While LiGaMD enables the sampling of multiple ligand unbinding events from the spike glycoprotein, obtaining a reliable PMF profile through post-LiGaMD analysis remains challenging. In this study, we conducted over 100 simulations, accumulating more than 50 μs of simulation time. However, only 8 complete ligand dissociation events were captured, resulting in a success rate of less than 8%. Given the low rate of capturing ligand unbinding, we had to increase the boost potential in LiGaMD to collect more data, which also compromises the reweighting accuracy. Consequently, we were unable to fully recover the original PMF profile due to the substantial variation in boost potential. Instead, we present the unweighted PMF of key reaction coordinates. While this unweighted PMF may not precisely depict the energetic changes of intermediates during ligand unbinding, it still represents the metastates within distinct unbinding pathways, as well as ligand-protein interactions at various stages.

Conformational selection and induced fit are two broadly recognized mechanisms in ligand binding and unbinding^54^. Conformational selection hypothesizes a protein with various conformational states, and ligand binding stabilizes one of these pre-existing states. Conversely, induced fit suggests a protein undergoes conformational changes upon ligand binding to optimize the interaction and create a complementary binding site. In our analysis of eight LA unbinding trajectories, we presented an even distribution of LA dissociation between Path A and Path B, which depended on two key factors: the gate distance between α1 and α3 and the position of the N343 glycan. Specifically, when the gate opens and the N343 glycan moves towards the RBD, LA dissociates along Path A; otherwise, it follows Path B. Regarding the gate distance, we noted a wide range of gate distances across various available spike protein structures, as outlined in Table S2, indicating the diversity of pre-existing gate distances within the spike. Moreover, our simulations of a holo spike glycoprotein revealed significant N343 glycan fluctuations throughout the simulation. During these fluctuations, the glycan could either point towards the adjacent RBM or the NTD. Hence, our findings strongly indicate that LA dissociation predominantly aligns with conformational selection principles.

The role of K417 in the LA dissociation is critical, as it forms polar interactions with LA during ligand unbinding. K417 engages in the spike-ACE2 binding exclusively in the SARS-CoV-2 system and has been suggested as contributing to the increased binding infinity of the SARS-CoV-2 spike to ACE2 as compared to the SARS-CoV spike^15,55^. When the RBD is in the “up” conformation, K417 is exposed to the solvent and forms polar interactions with residue D30 of ACE2 receptor. However, in the presence of LA bound to the spike, the RBD adopts a “down” conformation, resulting in K417 orienting towards the FFA binding pocket. This interaction suggests that LA binding contributes to stabilizing the RBD in the down conformation, consequently impeding the spike-ACE2 binding.

N-glycans are recognized for their impact on the spike glycoprotein. For example, N165 and N234 glycans have been found to enhance accessibility for spike-ACE2 binding^56,26^, while N282, N331, and N343 glycans provide shielding over the RBD in its down conformation^56,57,44^. Additionally, N616 and N1134 glycans have been identified as contributors to reduced spike glycoprotein stability^56,58^, while N717, N801, and N1074 glycans are associated with decreased viral infectivity^56,59^. Our study further elucidates the critical roles played by N122, N165, and N343 glycans in the process of ligand dissociation. When LA dissociates through Path A, interactions between N343 glycan and N122 and N165 glycans were found. When LA dissociates through Path B, N343 glycan interacts with a set of residues as depicted in Figure S7B-D, which are notably involved in the spike-ACE2 binding process. The presence of LA restrains the motion of the N165 glycan and the crucial residues that potentially form interactions with ACE2, suggesting that developing ligands with the capability to modulate glycan motions is a promising therapeutic strategy for addressing COVID-19 infection.

In our study, we found different LA dissociation behaviors in single-bound systems depending on the specific binding site. LAs bound to RBD_A_ exhibited rapid dissociation, requiring less applied boost in the simulation. Conversely, LA unbinding from RBD_B_ and RBD_C_ took longer and required higher boost levels. Moreover, compared to systems with a single bound ligand, those with multiple bound ligands demonstrated prolonged dissociation times and complex trajectories. These trajectories often involved multiple ligand flips and rebinding events throughout the unbinding process. Due to the limited number of available LA unbinding trajectories for analysis, the influence of the number of bound ligands on dissociation time and pathways remains unclear. Future efforts will focus on extensive sampling to collect enough successful unbinding events, enabling robust statistical analysis of ligand dissociation rates and a thorough exploration of how different ligand environments impact the dissociation process. Additionally, although the spike protein comprises three identical chains, glycosylation patterns vary across each chain. Investigating the influence of distinct glycan attachments on each chain will be crucial in understanding how different glycans contribute to preferred ligand dissociation pathways. Given current computational limitations, our simulations only encompassed the spike head. Future studies of the complete spike system, including the viral membrane and molecular crowding, will provide a biologically realistic environment for studying ligand diffusion. Expanding our investigation to include other ligands alongside LA will further enhance our understanding of ligand diffusion dynamics of the spike glycoprotein.

In summary, we applied LiGaMD simulations to reveal the dynamics of ligand dissociation from the SARS-CoV-2 spike glycoprotein. Through extensive simulations, we identified two distinct ligand dissociation pathways, underscoring the critical role of the gate formed by α1 and α3 helices in the RBD, as well as the glycan attached to N343, in determining the preferred dissociation routes. These crucial insights significantly enhance our understanding of the ligand unbinding process from the spike glycoprotein, with implications for drug discovery in the battle against COVID-19. The fundamental knowledge learned from this system can also be applied to the treatment of influenza or other viruses.

## Supporting Information

Supplemental information on the LiGaMD method and additional data can be found in the supporting document.

## Supporting information

Supplemental Tables and Figures and additional information on the LiGaMD method

## Acknowledgements

We express our gratitude to the National Energy Research Scientific Computing (NERSC) Center and the Wayne State University High-Performance Computing Center for supporting our LiGaMD simulations. This research is supported by the Wayne State University Start-up fund to Y.M.H.

