## Supplemental Tables and Figures and additional information on the LiGaMD method for "Mechanistic insights into ligand dissociation from the SARS-CoV-2 spike glycoprotein"

**Text S1: Ligand Gaussian accelerated molecular dynamics (LiGaMD)**

LiGaMD builds upon the Gaussian accelerated molecular dynamics (GaMD) method[1–3]. GaMD is an enhanced sampling technique that works by adding a harmonic boost potential to smooth the biomolecular potential energy surface and reduce energy barriers. Details of the GaMD method have been described in previous studies[1–3]. The ligand GaMD (LiGaMD) method has been developed for more efficient sampling of protein-ligand binding[4].

In the LiGaMD method, we consider a system composed of a ligand $L$ binding to a protein $P$ in a biological environment $E$. The system is comprised of $N$ atoms with positional coordinates $r \equiv \{\vec{r}_1, \cdots, \vec{r}_N\}$ and momenta $p \equiv \{\vec{p}_1, \cdots, \vec{p}_N\}$. The system Hamiltonian can be expressed as:

$$H(r,p) = K(p) + V(r), \tag{1}$$

where $K(p)$ and $V(r)$ represent the system's kinetic and total potential energies, respectively. Next, we can break down the potential energy into the following terms:

$$\begin{aligned} V(r) &= V_{P,b}(r_P) + V_{L,b}(r_L) + V_{E,b}(r_E) \\ &+ V_{PP,nb}(r_P) + V_{LL,nb}(r_L) + V_{EE,nb}(r_E) \\ &+ V_{PL,nb}(r_{PL}) + V_{PE,nb}(r_{PE}) + V_{LE,nb}(r_{LE}). \end{aligned} \tag{2}$$

where $V_{P,b}$, $V_{L,b}$ and $V_{E,b}$ are the bonded potential energies in protein $P$, ligand $L$ and environment $E$, respectively. $V_{PP,nb}$, $V_{LL,nb}$ and $V_{EE,nb}$ are the self nonbonded potential energies in protein $P$, ligand $L$ and environment $E$, respectively. $V_{PL,nb}$, $V_{PE,nb}$ and $V_{LE,nb}$ are the nonbonded interaction energies between $P$-$L$, $P$-$E$ and $L$-$E$, respectively. Classical molecular mechanics force fields calculate the nonbonded potential energies as[5–7]:

$$V_{nb} = V_{elec} + V_{vdW}. \tag{3}$$

where $V_{elec}$ and $V_{vdW}$ denote the electrostatic and van der Waals potential energies, respectively. Note that ligand binding only involves the nonbonded interaction energies of the ligand,

$V_{L,nb}(r) = V_{LL,nb}(r_L) + V_{PL,nb}(r_{PL}) + V_{LE,nb}(r_{LE})$. In LiGaMD, we add a boost potential selectively to the ligand nonbonded potential energy:

$$\Delta V_{L,nb}(r) = \begin{cases} \frac{1}{2} k_{L,nb} \left(E_{L,nb} - V_{L,nb}(r)\right)^2, & V_{L,nb}(r) < E_{L,nb} \\ 0, & V_{L,nb}(r) \geq E_{L,nb} \end{cases} \quad (4)$$

where $E_{L,nb}$ is the threshold energy for applying boost potential and $k_{L,nb}$ is the harmonic constant. The LiGaMD simulation parameters are derived in the same manner as in the GaMD algorithm[3]. When $E$ is set to the lower bound as the system maximum potential energy ($E = V_{max}$), the effective harmonic force constant $k_0$ can be calculated as:

$$k_0 = \min(1.0, k_0') = \min\left(1.0, \frac{\sigma_0}{\sigma_V} \cdot \frac{V_{max} - V_{min}}{V_{max} - V_{avg}}\right) \quad (5)$$

where $V_{max}$, $V_{min}$, $V_{avg}$, and $\sigma_V$ are the maximum, minimum, average and standard deviation of the boosted system potential energy, and $\sigma_0$ is the user-specified upper limit of the standard deviation of $\Delta V$ (e.g., 10 $k_B T$) for proper reweighting. The harmonic constant is calculated as $k = k_0 \cdot \frac{1}{V_{max} - V_{min}}$ with $0 < k_0 \leq 1$. Alternatively, when the threshold energy $E$ is set to its upper bound $E = V_{min} + \frac{1}{k}$, $k_0$ is set to:

$$k_0 = k_0'' \equiv \left(1 - \frac{\sigma_0}{\sigma_V}\right) \frac{V_{max} - V_{min}}{V_{avg} - V_{min}} \quad (6)$$

if $k_0$ is within the range of 0 to 1. Otherwise, $k_0$ is recalculated using Eq 5.

In addition to selectively boosting the bound ligand to accelerate its dissociation, an additional boost potential can be applied to unbound ligands, proteins, and solvent molecules, which can enhance the ligand rebinding to the protein. The second boost potential is determined using the system overall potential energy, excluding the nonbonded potential energy of the bound ligand:

$$\Delta V_D(r) = \begin{cases} \frac{1}{2} k_D \left(E_D - V_D(r)\right)^2, & V_D(r) < E_D \\ 0, & V_D(r) \geq E_D \end{cases} \tag{7}$$

where $E_D$ and $k_D$ are the corresponding threshold energy for applying the second boost potential and the harmonic constant, respectively. This leads to dual-boost LiGaMD with the total boost potential $\Delta V(r) = \Delta V_{L,nb}(r) + \Delta V_D(r)$. In this case, the strength of the individual boost potential term is determined by the user-defined threshold energies ($E_{L,nb}$ and $E_D$) and upper limits ($\sigma_{L,nb}$ and $\sigma_D$), the latter which determines the harmonic constants ($k_{L,nb}$ and $k_D$).

**Text S2: LiGaMD simulation input parameters**

An example of input parameters used in dual-boost LiGaMD simulations includes the following. The threshold energy is set to the upper bound for the ligand nonbonded boost potential and for the boost potential applied to the rest of the system. The user-specified upper limit of the boost potential is set to 6.0 kcal/mol for the ligand nonbonded term and 150.0 kcal/mol for the rest of the potential terms. The ligand nonbonded boost potential is applied to the ligand with residue number 3761.

Preparation Run:

***igamd = 7, irest_gamd = 0,***

***ntcmd = 2000000, nteb = 40000000, ntave = 500000,***

***ntcmdprep = 500000, ntebprep = 500000,***

***sigma0P = 6.0, sigma0D = 150.0, iEP = 2, iED = 2,***

***icfe = 1, ifsc = 1, gti_cpu_output = 0, gti_add_sc = 1,***

***timask1 = ':3761',***

***scmask1 = ':3761',***

| Chain | Site | Type | Structure | Sequence |
|---|---|---|---|---|
| A | N17 | FA2 | 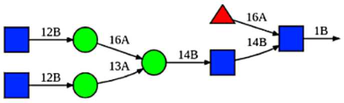 | bDGlcNAc(1→2)aDMan(1→6)[bDGlcNAc(1→2)aDMan(1→3)] bDMan(1→4)bDGlcNAc(1→4)[aLFuc(1→6)]bDGlcNAc(1→)PROA-17 |
| A | N61 | M5 | 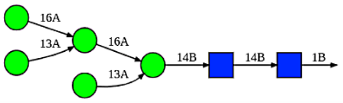 | aDMan(1→6)[aDMan(1→3)]aDMan(1→6) [aDMan(1→3)]bDMan(1→4)bDGlcNAc(1→4)bDGlcNAc(1→)PROA-61 |
| A | N122 | M5 | 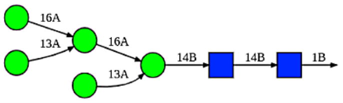 | aDMan(1→6)[aDMan(1→3)]aDMan(1→6) [aDMan(1→3)]bDMan(1→4)bDGlcNAc(1→4)bDGlcNAc(1→)PROA-122 |
| A | N165 | FA2G2S2 | 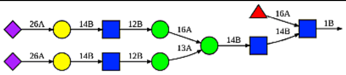 | xaDNeu5Ac(2→6)bDGal(1→4)bDGlcNAc(1→2)aDMan(1→6) [aDNeu5Ac(2→6)bDGal(1→4)bDGlcNAc(1→2)aDMan(1→3)] bDMan(1→4)bDGlcNAc(1→4)[aLFuc(1→6)]bDGlcNAc(1→)PROA-165 |
| A | N234 | M8 | 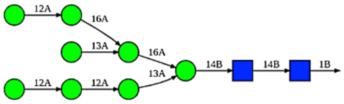 | aDMan(1→2)aDMan(1→6)[aDMan(1→3)]aDMan(1→6) [aDMan(1→2)aDMan(1→2)aDMan(1→3)] bDMan(1→4)bDGlcNAc(1→4)bDGlcNAc(1→)PROA-234 |
| A | N282 | FA3 | 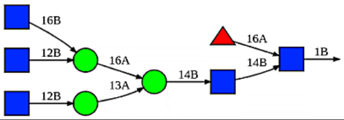 | bDGlcNAc(1→6)[bDGlcNAc(1→2)]aDMan(1→6) [bDGlcNAc(1→2)aDMan(1→3)]bDMan(1→4)bDGlcNAc(1→4) [aLFuc(1→6)]bDGlcNAc(1→)PROA-282 |
| A | N331 | FA2 | 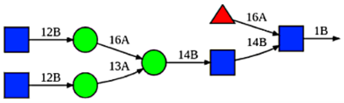 | bDGlcNAc(1→2)aDMan(1→6)[bDGlcNAc(1→2)aDMan(1→3)] bDMan(1→4)bDGlcNAc(1→4)[aLFuc(1→6)]bDGlcNAc(1→)PROA-331 |
| A | N343 | FA2 | 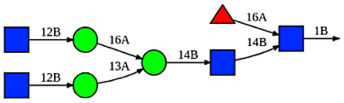 | bDGlcNAc(1→2)aDMan(1→6)[bDGlcNAc(1→2)aDMan(1→3)] bDMan(1→4)bDGlcNAc(1→4)[aLFuc(1→6)]bDGlcNAc(1→)PROA-343 |
| A | N616 | A2 | 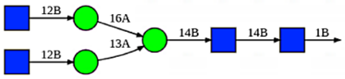 | bDGlcNAc(1→2)aDMan(1→6)[bDGlcNAc(1→2)aDMan(1→3)] bDMan(1→4)bDGlcNAc(1→4)bDGlcNAc(1→)PROA-616 |
| A | N709 | M6 | 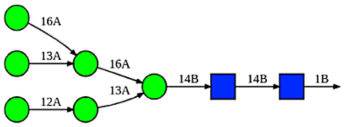 | aDMan(1→6)[aDMan(1→3)]aDMan(1→6)[aDMan(1→2)aDMan(1→3)] bDMan(1→4)bDGlcNAc(1→4)bDGlcNAc(1→)PROA-709 |
| A | N717 | Hybrid G1 | 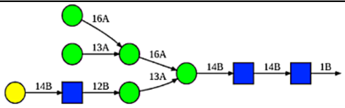 | bDGal(1→4)bDGlcNAc(1→2)aDMan(1→3)[aDMan(1→6)[aDMan(1→3)] aDMan(1→6)]bDMan(1→4)bDGlcNAc(1→4)bDGlcNAc(1→)PROA-717 |
| A | N801 | M6 | 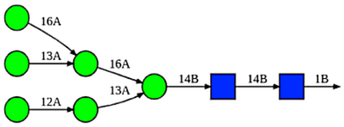 | aDMan(1→6)[aDMan(1→3)]aDMan(1→6)[aDMan(1→2)aDMan(1→3)] bDMan(1→4)bDGlcNAc(1→4)bDGlcNAc(1→)PROA-801 |
| A | N1074 | FA2G2S1 | 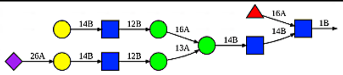 | aDNeu5Ac(2→6)bDGal(1→4)bDGlcNAc(1→2)aDMan(1→3) [bDGal(1→4)bDGlcNAc(1→2)aDMan(1→6)]bDMan(1→4)bDGlcNAc(1 →4) [aLFuc(1→6)]bDGlcNAc(1→)PROA-1074 |
| A | N1098 | FA2 | 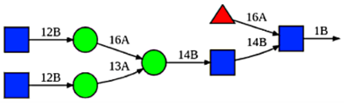 | bDGlcNAc(1→2)aDMan(1→6)[bDGlcNAc(1→2)aDMan(1→3)] bDMan(1→4)bDGlcNAc(1→4)[aLFuc(1→6)]bDGlcNAc(1→)PROA-1098 |
| A | N1134 | FA1 | 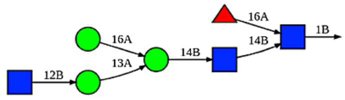 | bDGlcNAc(1→2)aDMan(1→3)[aDMan(1→6)]bDMan(1→4)bDGlcNAc(1 →4) [aLFuc(1→6)]bDGlcNAc(1→)PROA-1134 |

| | | | | |
|---|---|---|---|---|
| B | N17 | FA3 | 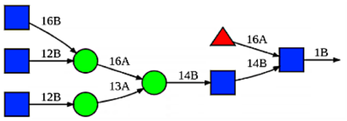 | bDGlcNAc(1→6)[bDGlcNAc(1→2)]aDMan(1→6)<br>[bDGlcNAc(1→2)aDMan(1→3)]bDMan(1→4)bDGlcNAc(1→4)<br>[aLFuc(1→6)]bDGlcNAc(1→)PROB-17 |
| B | N61 | M5 | 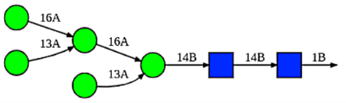 | aDMan(1→6)[aDMan(1→3)]aDMan(1→6)<br>[aDMan(1→3)]bDMan(1→4)bDGlcNAc(1→4)bDGlcNAc(1→)PROB-61 |
| B | N122 | FA2 | 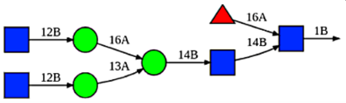 | bDGlcNAc(1→2)aDMan(1→6)[bDGlcNAc(1→2)aDMan(1→3)]<br>bDMan(1→4)bDGlcNAc(1→4)[aLFuc(1→6)]bDGlcNAc(1→)PROB-122 |
| B | N165 | M5 | 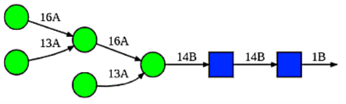 | aDMan(1→6)[aDMan(1→3)]aDMan(1→6)[aDMan(1→3)]<br>bDMan(1→4)bDGlcNAc(1→4)bDGlcNAc(1→)PROB-165 |
| B | N234 | M9 | 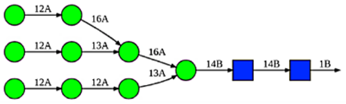 | aDMan(1→2)aDMan(1→6)[aDMan(1→2)aDMan(1→3)]aDMan(1→6)<br>[aDMan(1→2)aDMan(1→2)aDMan(1→3)]<br>bDMan(1→4)bDGlcNAc(1→4)bDGlcNAc(1→)PROB-234 |
| B | N282 | FA3G3S1 | 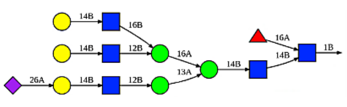 | aDNeu5Ac(2→6)bDGal(1→4)bDGlcNAc(1→2)aDMan(1→3)<br>[bDGal(1→4)bDGlcNAc(1→6)[bDGal(1→4)bDGlcNAc(1→2)]<br>aDMan(1→6)]bDMan(1→4)bDGlcNAc(1→4)<br>[aLFuc(1→6)]bDGlcNAc(1→)PROB-282 |
| B | N331 | FA2 | 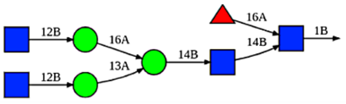 | bDGlcNAc(1→2)aDMan(1→6)[bDGlcNAc(1→2)aDMan(1→3)]<br>bDMan(1→4)bDGlcNAc(1→4)[aLFuc(1→6)]bDGlcNAc(1→)PROB-331 |
| B | N343 | FA1 | 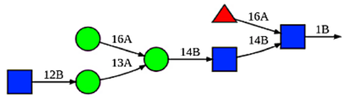 | bDGlcNAc(1→2)aDMan(1→3)[aDMan(1→6)]bDMan(1→4)bDGlcNAc(1<br>→4)<br>[aLFuc(1→6)]bDGlcNAc(1→)PROB-343 |
| B | N616 | FA2 | 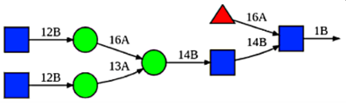 | bDGlcNAc(1→2)aDMan(1→6)[bDGlcNAc(1→2)aDMan(1→3)]<br>bDMan(1→4)bDGlcNAc(1→4)[aLFuc(1→6)]bDGlcNAc(1→)PROB-616 |
| B | N709 | M5 | 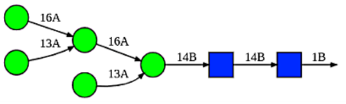 | aDMan(1→6)[aDMan(1→3)]aDMan(1→6)[aDMan(1→3)]<br>bDMan(1→4)bDGlcNAc(1→4)bDGlcNAc(1→)PROB-709 |
| B | N717 | M5 | 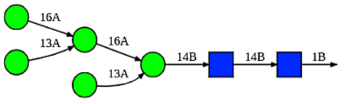 | aDMan(1→6)[aDMan(1→3)]aDMan(1→6)[aDMan(1→3)]<br>bDMan(1→4)bDGlcNAc(1→4)bDGlcNAc(1→)PROB-717 |
| B | N801 | M7 | 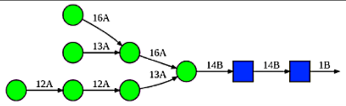 | aDMan(1→2)aDMan(1→2)aDMan(1→3)[aDMan(1→6)[aDMan(1→3)]<br>aDMan(1→6)]bDMan(1→4)bDGlcNAc(1→4)bDGlcNAc(1→)PROB-801 |
| B | N1074 | M5 | 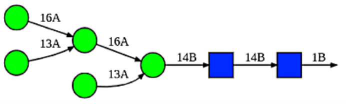 | aDMan(1→6)[aDMan(1→3)]aDMan(1→6)[aDMan(1→3)]<br>bDMan(1→4)bDGlcNAc(1→4)bDGlcNAc(1→)PROB-1074 |
| B | N1098 | A2 | 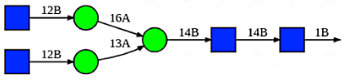 | bDGlcNAc(1→2)aDMan(1→6)[bDGlcNAc(1→2)aDMan(1→3)]<br>bDMan(1→4)bDGlcNAc(1→4)bDGlcNAc(1→)PROB-1098 |
| B | N1134 | FA3 | 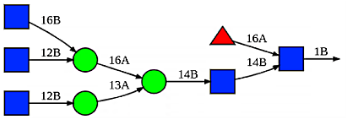 | bDGlcNAc(1→6)[bDGlcNAc(1→2)]aDMan(1→6)<br>[bDGlcNAc(1→2)aDMan(1→3)]bDMan(1→4)bDGlcNAc(1→4)<br>[aLFuc(1→6)]bDGlcNAc(1→)PROB-1134 |

| | | | | |
|---|---|---|---|---|
| C | N17 | FA3 | 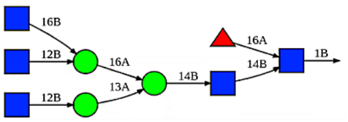 | bDGlcNAc(1→6)[bDGlcNAc(1→2)]aDMan(1→6) [bDGlcNAc(1→2)aDMan(1→3)]bDMan(1→4)bDGlcNAc(1→4) [aLFuc(1→6)]bDGlcNAc(1→)PROC-17 |
| C | N61 | M5 | 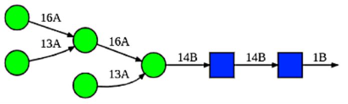 | aDMan(1→6)[aDMan(1→3)]aDMan(1→6) [aDMan(1→3)] bDMan(1→4)bDGlcNAc(1→4)bDGlcNAc(1→)PROC-61 |
| C | N122 | M5 | 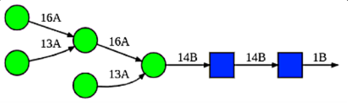 | aDMan(1→6)[aDMan(1→3)]aDMan(1→6) [aDMan(1→3)] bDMan(1→4)bDGlcNAc(1→4)bDGlcNAc(1→)PROC-122 |
| C | N165 | FA2G2S1 | 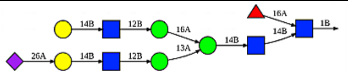 | aDNeu5Ac(2→6)bDGal(1→4)bDGlcNAc(1→2)aDMan(1→6) [bDGal(1→4)bDGlcNAc(1→2)aDMan(1→3)]bDMan(1→4)bDGlcNAc(1 →4) [aLFuc(1→6)]bDGlcNAc(1→)PROC-165 |
| C | N234 | M9 | 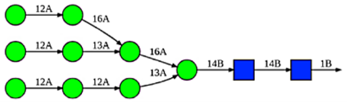 | aDMan(1→2)aDMan(1→6)[aDMan(1→2)aDMan(1→3)]aDMan(1→6) [aDMan(1→2)aDMan(1→2)aDMan(1→3)] bDMan(1→4)bDGlcNAc(1→4)bDGlcNAc(1→)PROC-234 |
| C | N282 | A2 | 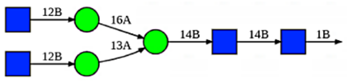 | bDGlcNAc(1→2)aDMan(1→6)[bDGlcNAc(1→2)aDMan(1→3)] bDMan(1→4)bDGlcNAc(1→4)bDGlcNAc(1→)PROC-282 |
| C | N331 | FA3G3S1 | 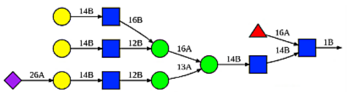 | aDNeu5Ac(2→6)bDGal(1→4)bDGlcNAc(1→2)aDMan(1→3) [bDGal(1→4)bDGlcNAc(1→6)[bDGal(1→4)bDGlcNAc(1→2)] aDMan(1→6)]bDMan(1→4)bDGlcNAc(1→4) [aLFuc(1→6)]bDGlcNAc(1→)PROC-331 |
| C | N343 | FA2 | 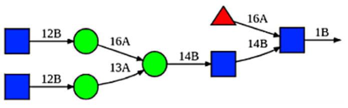 | bDGlcNAc(1→2)aDMan(1→6)[bDGlcNAc(1→2)aDMan(1→3)] bDMan(1→4)bDGlcNAc(1→4) [aLFuc(1→6)]bDGlcNAc(1→)PROC-343 |
| C | N616 | FA2 | 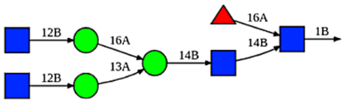 | bDGlcNAc(1→2)aDMan(1→6)[bDGlcNAc(1→2)aDMan(1→3)] bDMan(1→4)bDGlcNAc(1→4)[aLFuc(1→6)]bDGlcNAc(1→)PROC-616 |
| C | N709 | M5 | 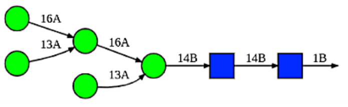 | aDMan(1→6)[aDMan(1→3)]aDMan(1→6) [aDMan(1→3)] bDMan(1→4)bDGlcNAc(1→4)bDGlcNAc(1→)PROC-709 |
| C | N717 | M6 | 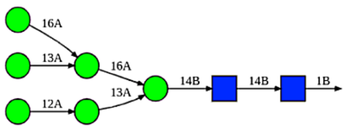 | aDMan(1→6)[aDMan(1→3)]aDMan(1→6)[aDMan(1→2)aDMan(1→3)] bDMan(1→4)bDGlcNAc(1→4)bDGlcNAc(1→)PROC-717 |
| C | N801 | M5 | 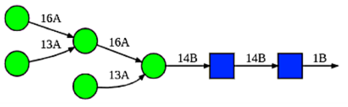 | aDMan(1→6)[aDMan(1→3)]aDMan(1→6) [aDMan(1→3)] bDMan(1→4)bDGlcNAc(1→4)bDGlcNAc(1→)PROC-801 |
| C | N1074 | M5 | 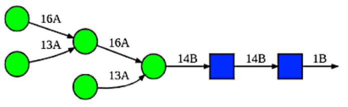 | aDMan(1→6)[aDMan(1→3)]aDMan(1→6) [aDMan(1→3)] bDMan(1→4)bDGlcNAc(1→4)bDGlcNAc(1→)PROC-1074 |
| C | N1098 | Hybrid G1S1 | 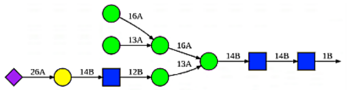 | aDNeu5Ac(2→6)bDGal(1→4)bDGlcNAc(1→2)aDMan(1→3) [aDMan(1→6)[aDMan(1→3)]aDMan(1→6)] bDMan(1→4)bDGlcNAc(1→4)bDGlcNAc(1→)PROC-1098 |
| C | N1134 | FA2 | 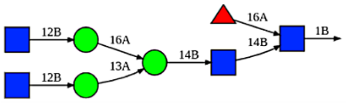 | bDGlcNAc(1→2)aDMan(1→6)[bDGlcNAc(1→2)aDMan(1→3)] bDMan(1→4)bDGlcNAc(1→4)[aLFuc(1→6)]bDGlcNAc(1→)PROC-1134 |

**Table S1:** Glycan attachments on the spike glycoprotein.

| PDB ID | Receptor-binding Domain | Ligand | Conformation | E340-A372 Distance |
|---|---|---|---|---|
| 6ZB5[8] | $RBD_A$ | Linoleate | Down | 13.31 |
| | $RBD_B$ | Linoleate | Down | 13.31 |
| | $RBD_C$ | Linoleate | Down | 13.31 |
| 6VXX[9] | $RBD_A$ | N/A | Down | 18.21 |
| | $RBD_B$ | N/A | Down | 18.21 |
| | $RBD_C$ | N/A | Down | 18.21 |
| 6VSB[10] | $RBD_A$ | N/A | Up | 18.27 |
| | $RBD_B$ | N/A | Down | 18.25 |
| | $RBD_C$ | N/A | Down | 17.14 |
| 6VYB[9] | $RBD_A$ | N/A | Down | 18.00 |
| | $RBD_B$ | N/A | Up | 18.38 |
| | $RBD_C$ | N/A | Down | 18.03 |
| 6ZGG[11] | $RBD_A$ | N/A | Down | 13.57 |
| | $RBD_B$ | N/A | Up | 11.93 |
| | $RBD_C$ | N/A | Down | 12.10 |
| 6ZGH[11] | $RBD_A$ | N/A | Down | 11.69 |
| | $RBD_B$ | N/A | Down | 12.43 |
| 7CAB[12] | $RBD_A$ | N/A | Down | 14.91 |
| | $RBD_B$ | N/A | Down | 14.91 |
| | $RBD_C$ | N/A | Down | 14.91 |
| 7DK3[13] | $RBD_A$ | N/A | Down | 16.67 |
| | $RBD_B$ | N/A | Down | 17.33 |
| | $RBD_C$ | N/A | Up | 15.20 |
| 7WZ1[14] | $RBD_A$ | N/A | Down | 15.55 |
| | $RBD_B$ | N/A | Up | 12.4 |
| | $RBD_C$ | N/A | Down | 17.72 |
| 7WZ2[14] | $RBD_A$ | N/A | Down | 18.97 |
| | $RBD_B$ | N/A | Up | 17.79 |
| | $RBD_C$ | N/A | Down | 18.63 |
| 6VW1[15] | $RBD_A$ | N/A | N/A | 19.37 |
| | $RBD_B$ | N/A | N/A | 19.20 |
| 7E3J[16] | $RBD_A$ | N/A | N/A | 19.75 |
| 7U0N[17] | $RBD_A$ | N/A | N/A | 19.27 |

| | | | | |
|---|---|---|---|---|
| | RBD$_B$ | ACE2 | N/A | 19.10 |
| 7W8S[18] | RBD$_A$ | ACE2 | N/A | 18.81 |
| 7WA1[18] | RBD$_A$ | ACE2 | N/A | 19.00 |
| 7C8D[19] | RBD$_A$ | ACE2 | N/A | 19.38 |
| 7C8J[20] | RBD$_A$ | ACE2 | N/A | 18.64 |
| 7CAH[12] | RBD$_A$ | H014 Fab | N/A | 17.29 |
| 7LM8[21] | RBD$_A$ | CV38-142 and COVA1-16 Fabs | N/A | 18.83 |
| 7LM9[21] | RBD$_A$ | CV38-142 Fab | N/A | 19.37 |
| 7TB8[22] | RBD$_A$ | A19-61.1 antibody | Up | 15.69 |
| | RBD$_B$ | A19-61.1 antibody | Down | 16.43 |
| | RBD$_C$ | B1-182.1 antibody | Up | 15.89 |
| 7TBF[22] | RBD$_A$ | B1-182.1 and A19-61.1 antibodies | N/A | 15.48 |
| 7WCD[23] | RBD$_A$ | TAU-2212 antibody | Down | 17.45 |
| | RBD$_B$ | TAU-2212 antibody | Down | 17.44 |
| | RBD$_C$ | TAU-2212 antibody | Down | 17.45 |
| 7WOG[14] | RBD$_A$ | 553-49 antibody | N/A | 16.59 |
| 7X7O[24] | RBD$_A$ | UT28K Fab | N/A | 14.95 |
| 8C8P[25] | RBD$_A$ | 10D12 antibody | N/A | 19.07 |

**Table S2:** Structure list of receptor-binding domains (RBDs) of the spike protein obtained from Protein Data Bank (PDB). The C$_\alpha$ distance between E340 and A372 was measured. The RBDs that cannot be determined either up or down conformation are noted as N/A.

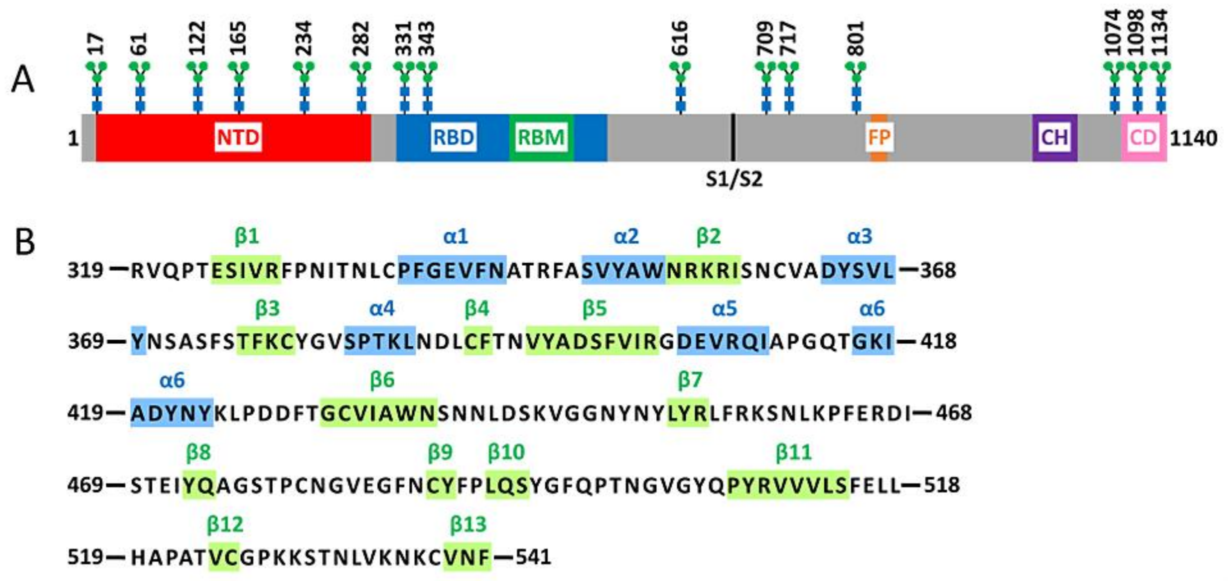


**Figure S1:** The structures and sequence of the SARS-CoV-2 spike glycoprotein. (A) Schematic of the spike protein primary structure for one chain: N-terminal domain (NTD, 16−291), receptor-binding domain (RBD, 319−541), receptor-binding motif (RBM, 438-506), furin cleavage site (S1/S2), fusion peptide (FP, 817−834), central helix (CH, 987−1034), connecting domain (CD, 1080−1140). Representative icons indicate N-glycans (blue and green) at N17, N61, N122, N165, N234, N282, N331, N343, N616, N709, N717, N801, N1074, N1098, and N1134. (B) Sequence and secondary structures of the spike RBD. Blue and green indicate α helices and β sheets, respectively.

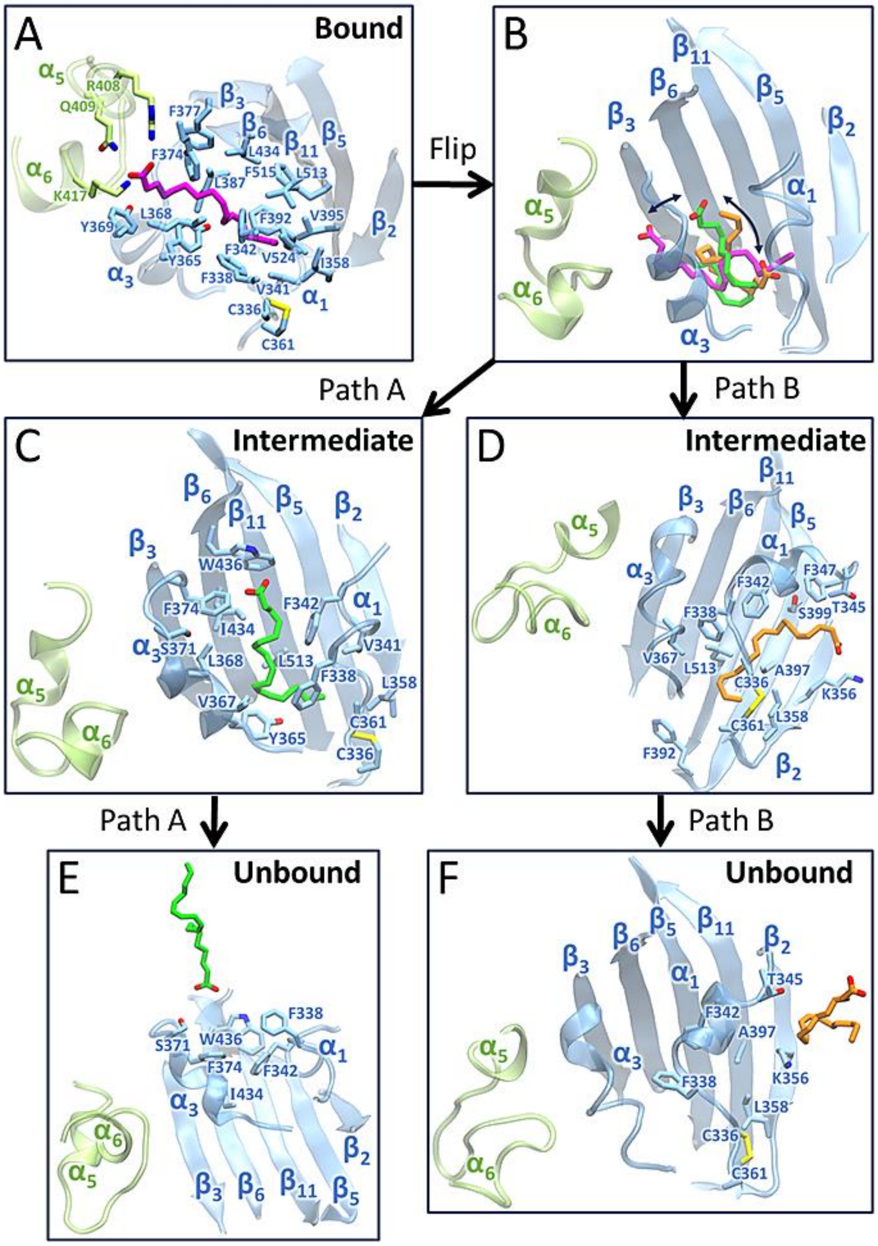


**Figure S2:** Movement of LA along two dissociation pathways. (A) LA is tightly bound in the FFA binding pocket. (B) LA undergoes a flipping motion, initiating the dissociation process. (C) LA moves along Path A. (D) LA moves along Path B. (E) LA fully dissociates from the binding pocket along Path A. (F) LA fully dissociates from the binding pocket along Path B.

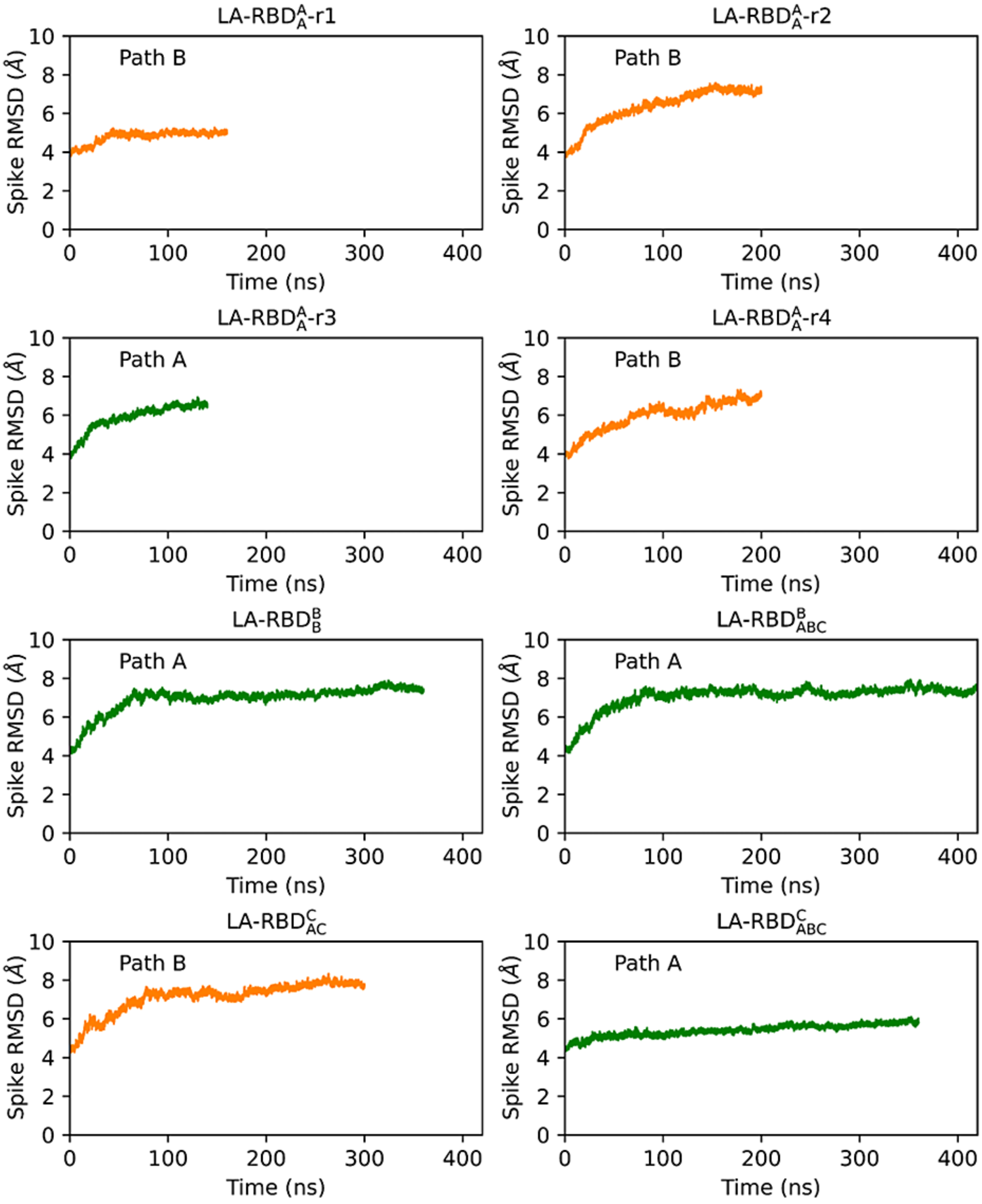


**Figure S3:** RMSD analysis of the full spike protein from eight dissociation trajectories. Green and orange indicate trajectories dissociating along Path A and B, respectively.

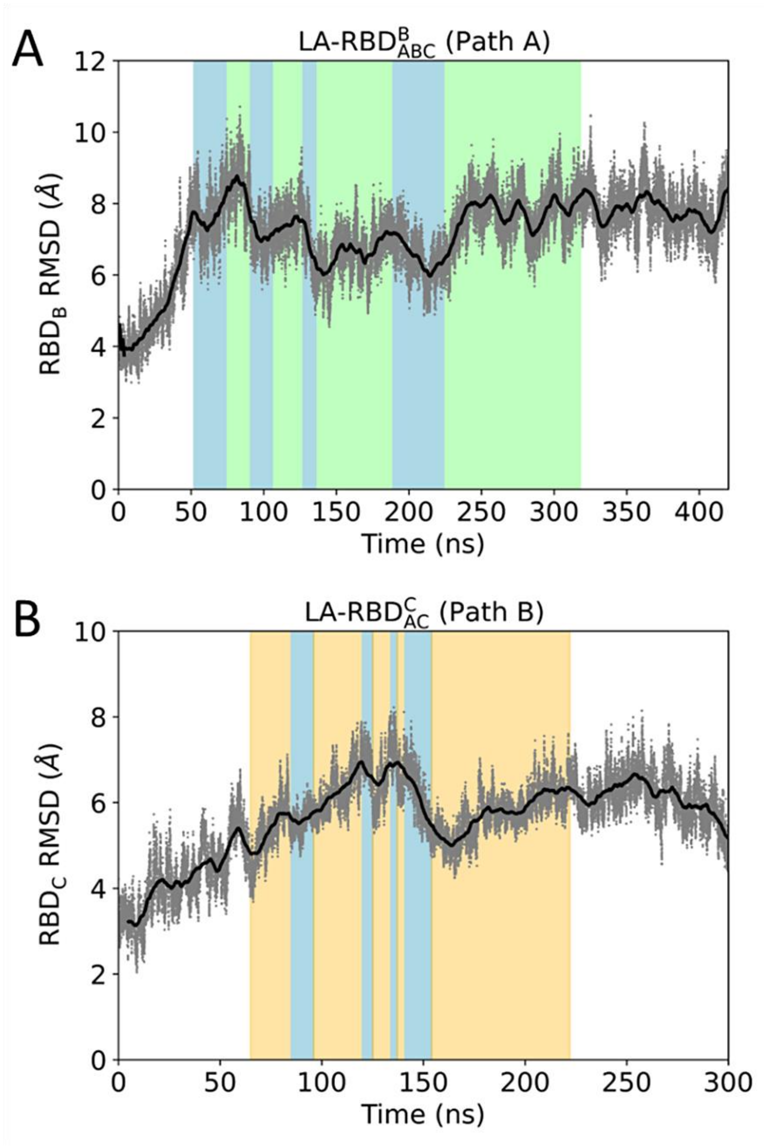


**Figure S4:** RMSD analysis of the spike RBD in complex with a boosted LA for trajectories, LA-RBD$^{B}_{ABC}$ (A) and LA-RBD$^{A}_{AC}$ (B). The color highlights different LA states: fully bound (blue), traveling along Path A (green), and moving along Path B (orange).

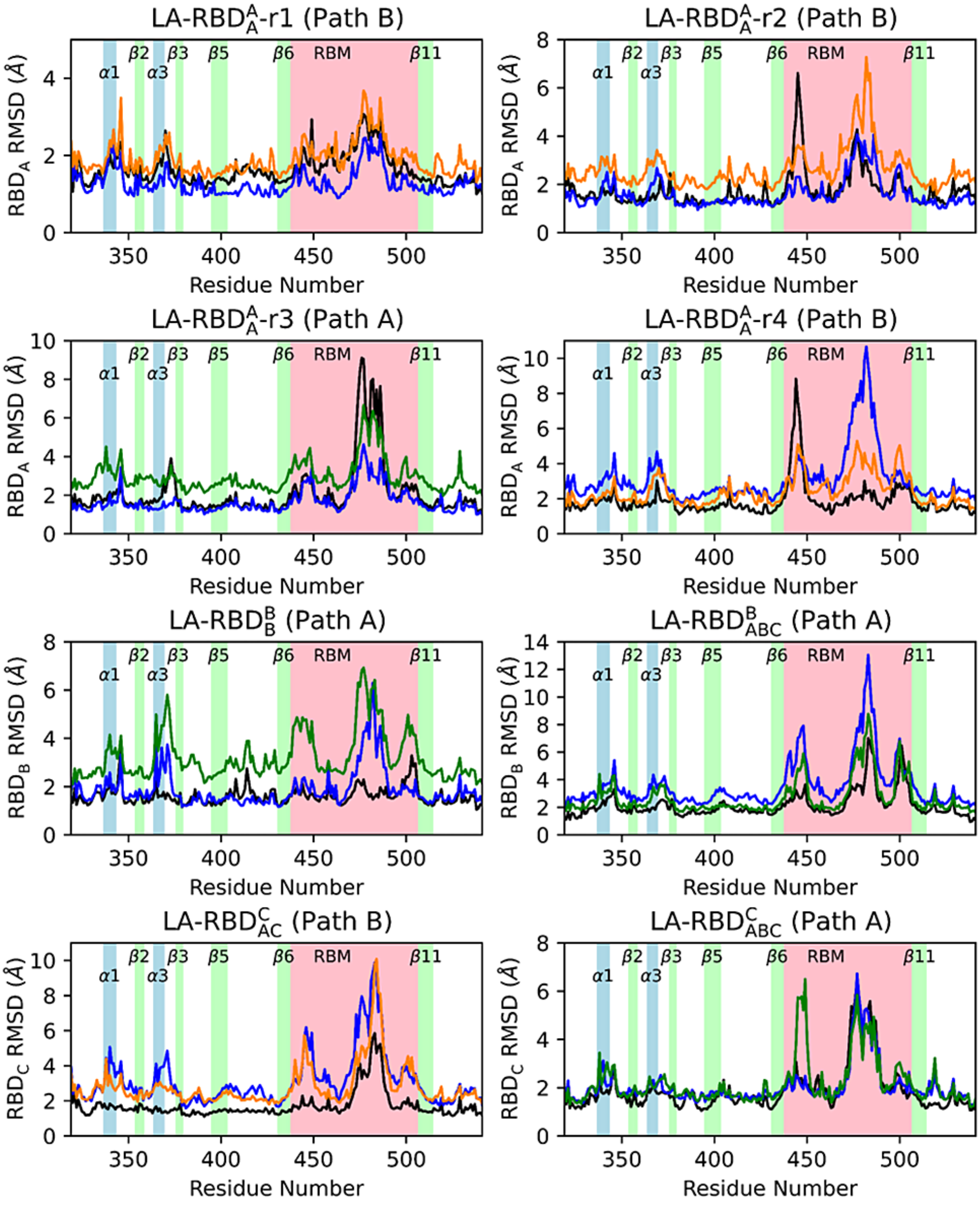
LA-RBD$^{A}_{A}$-r1 (Path B)
LA-RBD$^{A}_{A}$-r2 (Path B)
LA-RBD$^{A}_{A}$-r3 (Path A)
LA-RBD$^{A}_{A}$-r4 (Path B)
LA-RBD$^{B}_{B}$ (Path A)
LA-RBD$^{B}_{ABC}$ (Path A)
LA-RBD$^{C}_{AC}$ (Path B)
LA-RBD$^{C}_{ABC}$ (Path A)
$RBD_A$ RMSD (Å)
$RBD_B$ RMSD (Å)
$RBD_C$ RMSD (Å)
Residue Number
α1
α3
β2
β3
β5
β6
RBM
β11

**Figure S5:** RMSF analysis of the spike RBD in complex with LA from eight dissociation simulations. Line color represents RMSF profiles derived from trajectories during fully bound state (blue), traversal of Path A (green), traversal of Path B (orange), and after dissociation of the ligand (black). The highlighted colors correspond to different secondary structures: blue represents α1 (residues 337-343) and α3 (residues 364-369) helices, green corresponds to each β-strand (β2 residues 354-358, β3 residues 376-379, β5 residues 395-403, β6 residues 431-437, and β11 residues 507-514), and pink denotes the receptor-binding motif (RBM, residues 438 to 506).

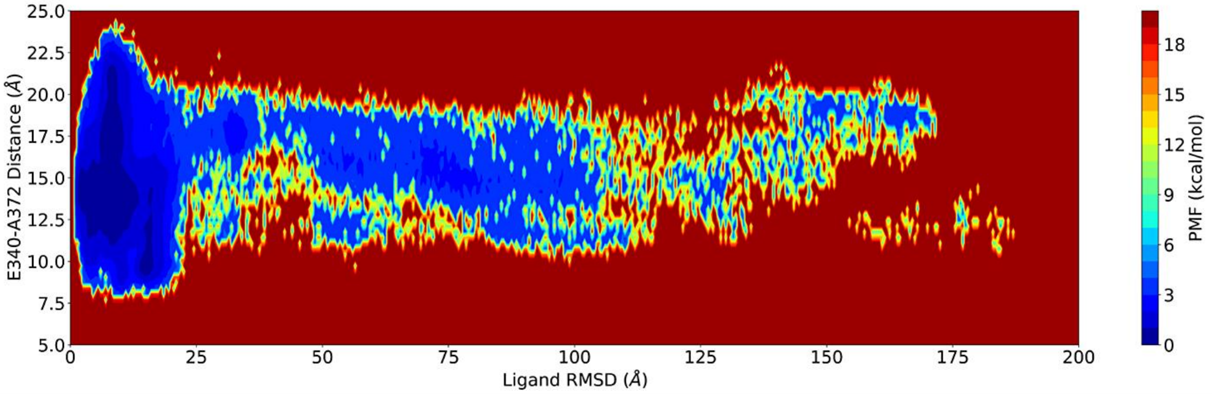


**Figure S6:** PMF profile of LA dissociation. The plot depicts the PMF changes with the ligand RMSD and the gate distance between E340 and A372, including the states after LA fully dissociates with the ligand RMSD exceeds 30 Å.

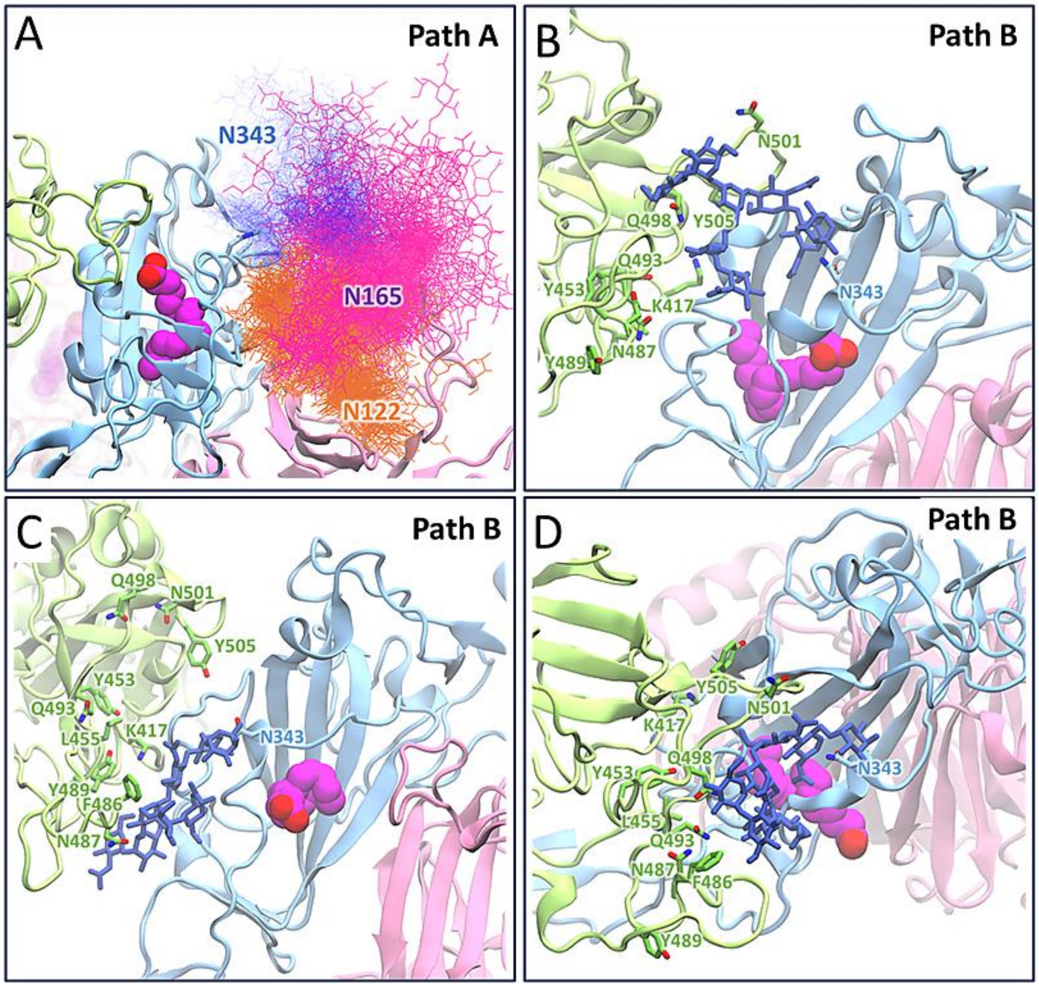


**Figure S7:** Movement of N343-glycan during LA dissociation. As LA dissociates from Chain A (blue) along Path A, the N343-glycan interacts with the glycans on N122 and N165 on the NTD in Chain B (pink) (A). Conversely, during LA dissociation along Path B, the glycan interacts with the residues of the RBM in Chain C (green), as illustrated in (B), (C), and (D).
